# SWI/SNF component BAF250a coordinates OCT4 and WNT signaling pathway to control cardiac lineage differentiation

**DOI:** 10.1101/524454

**Authors:** Ienglam Lei, Shuo Tian, Victor Chen, Yong Zhao, Zhong Wang

## Abstract

Dissecting epigenetic mechanisms controlling early cardiac differentiation will provide insights into heart regeneration and heart disease treatment. SWI/SNF complexes remodel nucleosomes to regulate gene expression and play a key role in organogenesis. Here we reported a unique function of BAF250a in regulating the physical interaction of OCT4 and β-CATENIN during cardiac lineage differentiation from human ESCs. BAF250a deletion greatly reduced the physical interaction between OCT4 and β-CATENIN but did not alter the expression of β-CATENIN and OCT4 in the mesodermal progenitor cells. BAF250a ablation led to decreased recruitment of OCT4 and β-CATENIN at promoters of key mesodermal lineage genes, such as MESP1 and EOMES. Subsequently, the expression of lineage specific genes was down-regulated whereas the expression of pluripotent genes was up-regulated. In parallel, BAF250a ablation also altered recruitments of OCT4 and β-CATENIN to the promoter of CCND2 and CCND3, two key genes for S phase entry during cell cycle. Consequently, BAF250a deletion led to prolonged S phase in Mesp1^+^ cardiac progenitor cells, which in turn inhibited efficient differentiation of Mesp1^+^ to Isl1^+^ cells. Furthermore, BAF250a deletion abolished the interaction of OCT4 and BRG1 in mesoderm, suggesting that BAF250a is the key component in SWI/SNF complex that determines the interaction of Oct4/β-catenin in mesoderm. In contrast, we found that BAF250a did not regulate the OCT4/β-CATENIN interaction during neuroectoderm differentiation. Altogether, our results suggest that BAF250a specifically controls proper cardiac mesoderm differentiation by reorganizing the binding of OCT4/β-CATENIN and regulates both key lineage differentiation genes and cell cycle genes coincided in response to WNT/β-CATENIN signal.

**Highlights:** BAF250a is required for hESC cardiac differentiation

BAF250a is required for the assembly of Brg1/OCT4/β-CATENIN complex and the recruitment of OCT4/β-CATENIN to cardiac genes

BAF250a is dispensable for the interaction of OCT4/β-CATENIN interaction in neuroectoderm differentiation

BAF250a interacts with OCT4/β-CATENIN to promote cardiac differentiation by regulating cell cycle.

## 1. Introduction

Dissecting the molecular mechanisms underlying the differentiation of pluripotent stem cells (PSC) into tissue specific progenitors and terminal differentiated cell types are key to understanding organ development and regeneration. Within heart, embryonic cardiac progenitor cells (CPCs) are promising cell sources for cell-based therapies for heart disease. Direct differentiation of CPCs from PSCs has been achieved based on the knowledge of embryonic development (1,2). Each step of differentiation is tightly regulated by stage-specific signals and epigenetic regulation(3). In particular, an initial WNT activation and subsequent WNT inhibition is required for CPC differentiation(4). In parallel, epigenetic regulations including histone modification and chromatin remodeling are also believed to contribute to the precise of cardiac differentiation(5). However, how WNT signaling and epigenetic regulation work together to direct cardiac differentiation is poorly understood.

SWI/SNF ATP-dependent chromatin remodeling complexes modulates the chromatin accessibility by reorganizing nucleosomes to regulate chromatin accessibility and are essential in organogenesis(6). BAF complexes also interact directly with other epigenetic factors and transcription factors in regulating gene expression. Studies from numerous research groups including us reveal that BAF complexes are important for embryonic stem cell (ESC) pluripotency and cardiogenesis(7–11). Nevertheless, the underlying epigenetic mechanisms of BAF mediated cardiac lineage specification are not well understood.

Cell cycle control is another key regulatory process during differentiation. Recent studies have identified cross-talk between cell cycle and cell fate determination of human ESCs (hESCs)(12,13). The hESCs have a unique cell cycle pattern with a short G1 phase, while their differentiation lengthened the G1 phase(14). The differentiation potential of hESCs vary at different cell cycle stage. It is shown that the activity of Activin/Nodal signal is regulated by cyclinD-CDK4/6 complex in early and late G1 stage(12). S and G2 phases have further been found to actively promote pluripotent state(15). These studies suggest that cell cycle phases could play an essential role in commitment of mesodermal cells into cardiac lineages, a notion that remains to be confirmed.

In this study, we define a unique function of BAF in regulating cardiac lineage differentiation in human ESCs by examining a key regulatory subunit BAF250a. By taking advantage of a conditional BAF250a knockout human ESCs we generated(16), we observed that BAF250a is required for the physical interaction of OCT4 and β-CATENIN in mesoderm cells. Consequently, BAF250a deletion affected the recruitment of OCT4 and β-CATENIN at target gene promoters and altered gene expression of differentiation, pluripotent, and cell cycle genes. We further revealed that the BAF250a deletion led to S phase population increase of mesodermal cells which have less differential potential into Isl1^+^ CPC cells. In addition, we found that BAF250a was dispensable for the interaction of OCT4 and β-CATENIN during neuroectoderm differentiation and BAF250a deficiency did not affect neuroectoderm differentiation. These results suggest an important role of BAF250a in reorganizing the binding of OCT4/β-CATENIN and regulating both key genes for lineage differentiation and cell cycle phases for proper cardiac mesoderm differentiation in response to WNT signal.

## 2. Methods

### 2.1. hESC culture and differentiation

hESCs were maintained on Matrigel-coated dishes with mTseR1 medium as described(34). The differentiation of hESC to cardiomyocyte was performed using sequential WNT modulation (4). Briefly, hESCs was dissociated into single cells with Versene solution. 1 million hESCs were plated to a 35mm dish with 10 μM Rock inhibitor. 6 μM GSK3 inhibitor CHIR99021 in chemical defined medium (CDM, 0.5% Human Albumin in RPMI1640) was added to the confluent hESC to induce the differentiation for two days. Cells were recovered for two days in CDM, then treated with 2 μM IWR1 in CDM for another two days. The culture medium was changed to CDM for two days, then 3% Knockout serum replacement was added to CDM in subsequent cultures. For neuroectoderm differentiation, the hESC monolayer culture was subject to differentiation medium (20% KOSR in KO DMEM) containing 500nM LDN193189 and 10 μM SB431542 for 4 days(26). Then the differentiation medium was changed to containing 25%, 50%, 75% and 100% N2 medium every other day from day 4 to day 12. The medium was changed every day. Day 12 cells were harversted for FACS and RT-qPCR analyses.

### 2.2. Western Blot

Cells were lysed in 0.25% Triton X-100; 50 mM Tris HCl, pH 7.4; 150 mM NaCl; 1 X protease inhibitors (Roche) then spun at 16000xg for 5 min at 4°C. Laemmli sample buffer was added before boiling for 5 minutes. Primary antibody OCT4 (sc-8628, 1:1000), BRG1(ab4081, 1:1000), BAF250a (sc-32761, 1:1000), T (sc-17745, 1:1000), active β-CATENIN (CST 4270, 1:1000) and β-ACTIN (CST4970, 1:2000) was applied to the blots at 4°C overnight. Secondary antibody from LiCOR was used for detection.

### 2.3. Co-immunoprecipitation

The co-immunoprecipitation was performed as previously described(8). Briefly, day 1 differentiated cells were harvested in cold PBS and extracted for 30 min at lysis buffer (20mM HEPES pH8.0, 150mM NaCl, 1% NP-40, 2mM EDTA) with proteinase inhibitors. After centrifugation, 5% of supernatant was kept in 4°C as input. The remaining supernatant was separated into equal volume and incubated with 1 μg of OCT4 or rabbit IgG with 20 μL Dynabeads at 4°C overnight. The complex was washed 4 times in wash buffer (20mM HEPES pH8.0, 150mM NaCl, 0.1% NP-40, 2mM EDTA). Beads were boiled for 5 minutes in 1x SDS buffer to denature proteins. After SDS-PAGE, western blots were performed using antibody against β-CATENIN (CST8480, 1:1000) and active β-CATENIN (1:1000).

### 2.4. Immunostaining

Embryos were fixed in 4% paraformaldehyde at 4°C overnight and embedded in paraffin. Sections were collected using microtome. Sections were blocked with 10% horse serum in PBS buffer containing 0.1% Tween 20 and 0.1% Triton X-100 for 1 h at room temperature and incubated with goat anti-T (sc-17745 1:100), mouse anti-OCT4(sc-8628, 1:200) and rabbit active β-CATENIN (CST 4270, 1:200) at 37°C for 1 h. Then sections were washed three times with PBS buffer with 0.1% Tween 20 before being incubated with a donkey anti-goat Alexa 488-conjugated, donkey anti-mouse Alexa 594-conjugated and donkey anti-rabbit Alexa 647-conjugated antibodies at room temperature for 1 h. After washes, section slides were mounted and analyzed by fluorescent microscopy.

### 2.5. Proximity ligation assay (PLA)

PLA was performed using Duolink^®^ In Situ Red kit (DUO92101) according to Sigma’s protocol. Briefly, E7 embryo sections were dewaxed with xylene, and rehydrate with descending alcohol. Antigen retrieval was performed by microwaving section in pH9 Tris-EDTA buffer for 15 minutes. The section was permeabilized with 0.2% Triton X-100 and blocked in 5% horse serum for 1 hour. Incubation with primary antibodies OCT4 and β-CATENIN were performed at 4°C overnight. Sections were washed for three times in PBST. The ligation and detection of OCT4/β-CATENIN by proximity probes was performed according the manufacture’s protocol.

### 2.6. Chromatin immunoprecipitation

ChIP experiments were performed as previously described(8). Briefly, cells were fixed in 1% formaldehyde for 10 minutes and quenched with 0.125M glycine for 5 minutes. Cells were then harvested and sonicated into 200-1000bp using branson sonifier. Chromatin solution was incubated with Dynabeads and antibodies against OCT4 or β-CATENIN overnight. Beads were washed four times with LiCl wash buffer followed by one wash of TE buffer then eluted with elution buffer at 65 °C. DNA was purified using phenol:chloroform extraction. Enrichment of immunoprecipitated DNA was then validated by quantitative PCR.

## 3. Results

### 3.1. BAF250a Deletion disrupted cardiomyocyte differentiation but not primitive induction of hESCs

To examine the role of BAF250a in human cardiac differentiation, we used direct differentiation protocol (Fig 1A) in a 4-hydroxytamoxifen(4-OHT)-inducible BAF250a knockout (KO) hESC line we generated(4,16). 4-OHT was added one day before induction of differentiation (day -1) and BAF250a was undetectable after 24 hours of 4-OHT treatment. The percentage of T+ cell was not affected after BAF250a deletion (Fig 1B). However, the T expression was slightly decreased after BAF250a deletion at day 1 of differentiation (Fig 1C). In contrast, deletion of BAF250a at ESC stage led to significant reduction of cTNT+ CMs (Fig 1D). We further found that BAF250a deletion in ESC led to decreased differentiation efficiency of MESP1+ cell and ISL1+ CPC cells (Fig 1D). These results indicated that BAF250a was dispensable in the primitive induction during hESC differentiation and was required for efficient cardiomyocyte differentiation from mesodermal progenitor cells.

**Figure 1.**
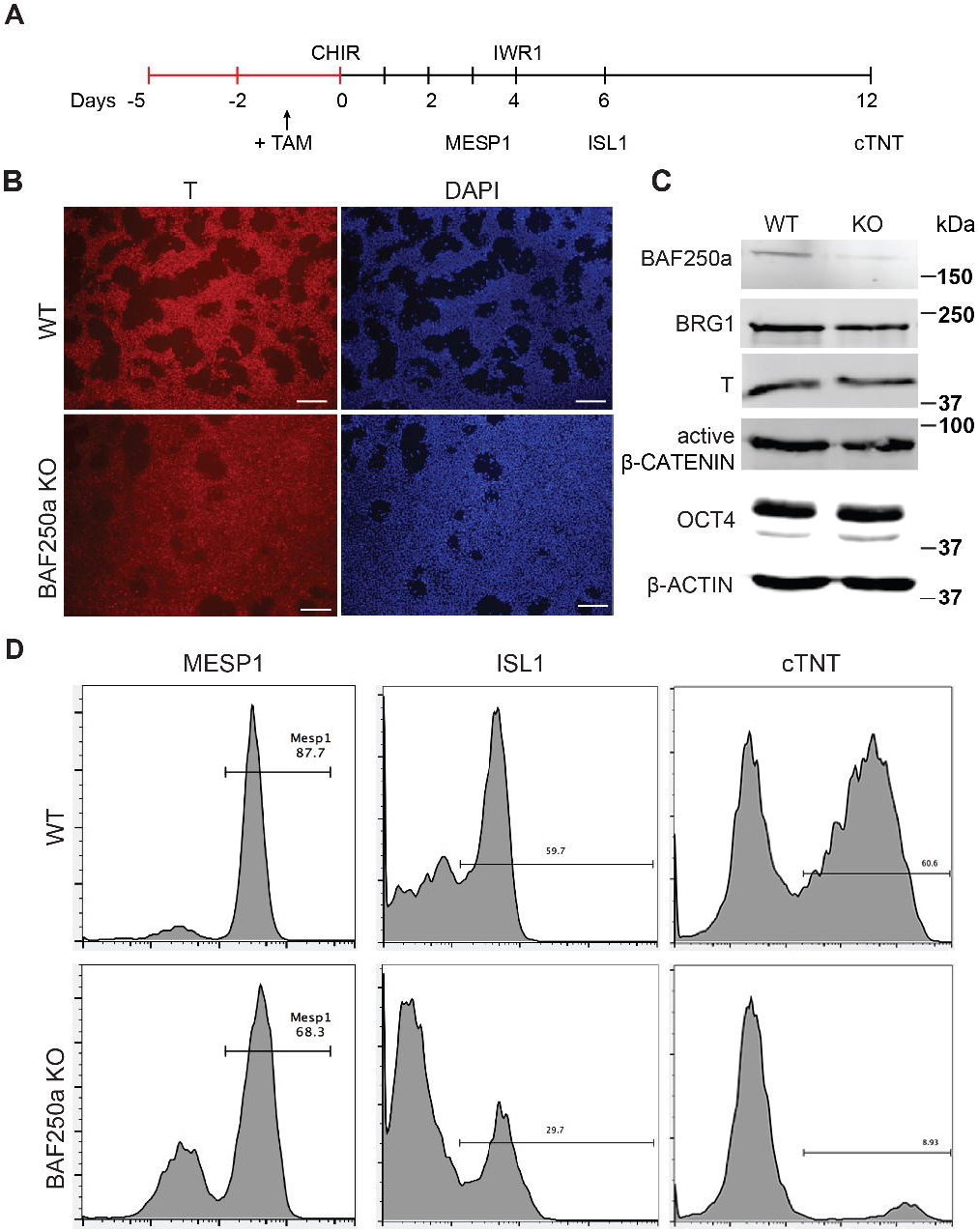
BAF250a deletion in hESCs led to cardiac differentiation defects. (A) Schematic diagram of cardiac differentiation. (B) Immunostaining of T at differentiation day 1 in both WT and BAF250a KO cells. (C) Western blot of BAF250a, BRG1, T, β-CATENIN and OCT4 in WT and BAF250a KO cells at differentiation day 1. β-actin was used as a loading control. (D) FACS analysis of differentiation of WT and BAF250a KO hESC to Mesoderm, cardiac progenitors, and cardiomyocytes using MESP1, ISL1 and cTNT antibodies at differentiation day 3, 6 and 10. Scale bar: 50μm

### 3.2. BAF250a was required for OCT4/β-CATENIN interaction

WNT/β-CATENIN signaling pathway is essential for gastrulation and mesodermal formation and are key factors in the established cardiac lineage differentiation from hESCs(17). We therefore examined the expression and activity of WNT/β-CATENIN with and without BAF250a during hESC differentiation. We found that the expression of active form β-CATENIN in day 1 differentiated cells was not changed after BAF250a deletion (Fig 1C). Moreover, the expression of OCT4, which has been shown to have important roles in mesoderm differentiation(18,19), was not affected by BAF250a deletion (Fig 1C). We then hypothesized that BAF250a might be required for the physical interaction of OCT4 and β-CATENIN in the mesoendoderm cells during hESC differentiation, as BAF250a interacts with OCT4 in ESCs(20) and OCT4/β-CATENIN interaction is important for proper mesoderm differentiation(19). Co-immunoprecipitation experiments at day 1 of differentiation demonstrated decreased interaction of OCT4 and β-CATENIN or active β-CATENIN in BAF250a depleted cells (Fig 2A, B), indicating that BAF250a was required for efficiency physical interaction of OCT4 and β-CATENIN. Consistent with this finding, we detected that OCT4 and β-CATENIN was co-localized at T+ cells at E7.0 mouse embryos (Fig 2C). Moreover, OCT4 and β-CATENIN interaction was also observed in E7.0 embryos by Proximal ligation assay (PLA) (21)(Fig 2D). As BRG1, the core catalytic subunit of SWI/SNF complex, has been shown to interact with OCT4 in ESCs (22) and BAF250a is considered a regulatory subunit and does not affect the formation of the core SWI/SNF complex, we wonder whether the observed BAF250a and OCT4/β-CATENIN interaction is secondary to BRG1 and OCT4/β-CATENIN interaction. Surprisingly, we found that the interaction of OCT4 and BRG1 was abolished in BAF250a KO cells (Fig 2E&F), indicating that BAF250a is a key subunit determining the interaction of SWI/SNF with OCT4 in mesoderm cells. Together with our previous finding that BAF250a is essential for mesoderm formation during mouse embryogenesis(23), our results suggested that BAF250a-mediated OCT4/β-CATENIN interaction is key to cardiac mesoderm specification during development.

**Figure 2.**
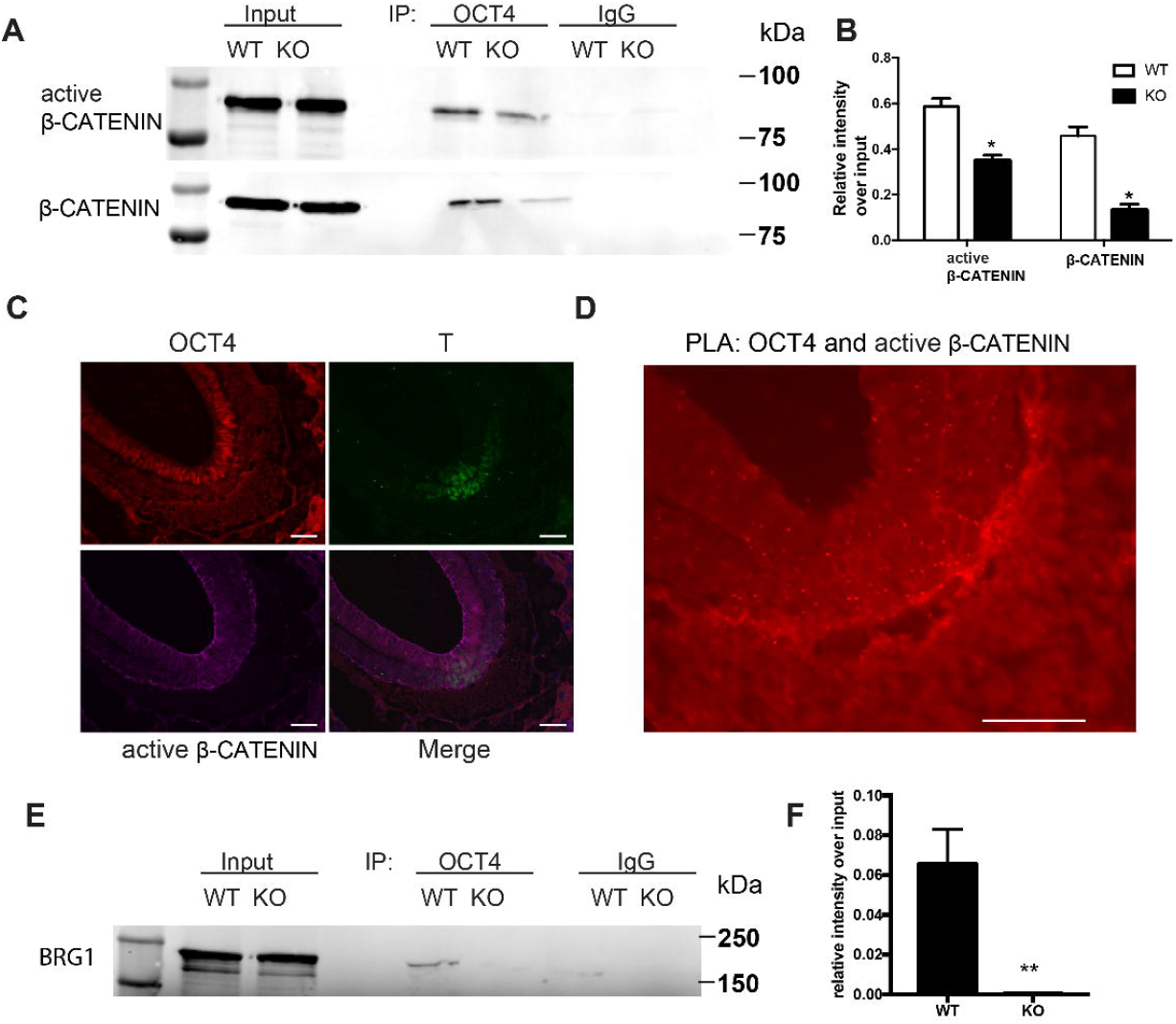
BAF250a regulated proper OCT4 and β-CATENIN interaction during differentiation. (A) Immunoprecipitation of day 1 differentiation mesodermal cells with anti-OCT4 antibody, followed by Western blotting to detect β-CATENIN and active β-CATENIN. 5% of cell lysates were used as input controls. (B) Relative band intensity of precipitated β-CATENIN and active β-CATENIN in WT and BAF250 KO cells. (C) Immunostaining of OCT4, T, and active β-CATENIN showing their colocalization at E7.0 embryos. (D) OCT4 and active β-CATENIN interaction in E7.0 embryos detected by PLA. Scale bar: 50μm (E) Immunoprecipitation of day 1 differentiation mesodermal cells using anti-OCT4 antibody, followed by Western blotting to detect BRG1. 5% of cell lysates were used as input controls. (F) Relative band intensity of precipitated BRG1 in WT and BAF250 KO cells. *p<0.05, **p <0.01, n=3.

### 3.3. Proper OCT4 and β-CATENIN interaction mediated by BAF250a guided cardiac mesoderm gene expression

To reveal the function BAF250a-mediated OCT4/β-CATENIN interaction in the differentiation of cardiac mesoderm, we first investigated the mesodermal gene expression at day 1 and day 3 after hESC differentiation. MESP1 and EOMES expression was significantly down regulated in BAF250a KO cells. We also found that the pluripotent gene SOX2 and NANOG expression were not properly turned down in BAF250a KO cells. In addition, we also detected upregulation of neural gene expression TFAP2A (Fig 3A). These results suggest that BAF250a is required for proper silencing of pluripotent genes and neural genes during differentiation. Given that SWI/SNF complex is essential to create accessible chromatins (24), we then examined the recruitment of OCT4 and β-CATENIN at MESP1, EOMES and TFAP2A promoters, as OCT4/ β-CATENIN interaction is important for the activation of cardiac mesoderm genes and repressing neural genes. ChIP assays showed decreased recruitment of OCT4 and β-CATENIN to MESP1 and EOMES promoters, whereas only decrease of OCT4 recruitment was found at TFAP2A promoters (Fig 3B, C). In addition, we found that the H3K27me3 modifications at MESP1 and EOMES promoters were not efficiently removed in BAF250a KO cells, while lower H3K27me3 level was found in TFAP2A promoters (Fig 3D). Considering that OCT4/β-CATENIN replace OCT4/Sox2 to promote cardiac mesoderm differentiation(25) and play a critical role in neural gene repression(19), it is likely that BAF250a is required for efficient OCT4/β-CATENIN recruitment to targets for both silencing pluripotent genes and activating cardiac mesoderm genes.

**Figure 3.**
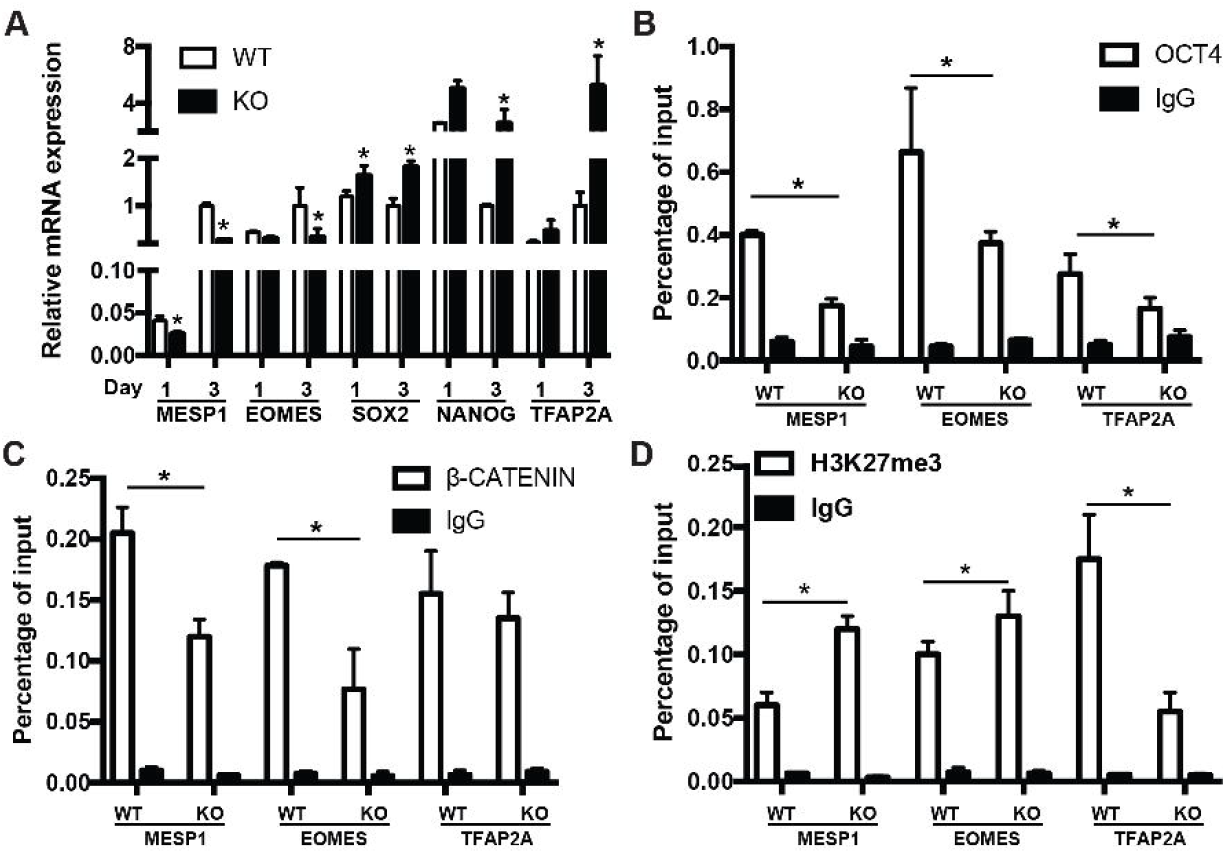
BAF250a controlled cardiac mesoderm gene expression. (A) Expression of MESP1, EOMES, SOX2, NANOG, and TFAP2A at differentiation day 1 and day 3. ChIP assays with (B) Anti-OCT4, (C) anti-β-CATENIN and (D) anti-H3K27me3 antibodies using day 1 differentiated cells at MESP1, EOMES and TFAP2A promoters. *p <0.05, n=3.

To examine whether the role of BAF250a is specific for mesoderm lineage, we next determined the differentiation efficiency of neuroectoderm from hESC using a dual inhibitor neuroectoderm differentiation assay (26) using both WT and BAF250a KO hESCs. We found that the percentage of Pax6+ neuroectoderm cells was not affected in BAF250a KO cells at days 12 after inducing differentiation (Fig 4A). The expression levels of TFAP2A and PAX6 were also comparable between WT and BAF250a KO cells (Fig 4B). Importantly, we found that the interaction of OCT4 and β-CATENIN was not affected in BAF250a KO cells at day 2 of neuroectorderm differentiation (Fig 4C&D). Furthermore, the recruitment of OCT4 and β-CATENIN to TFAP2A and PAX6 was not changed in BAF250a KO cells (Fig 4E&F). These results suggested that BAF250a was dispensable for the interaction of OCT4 and β-CATENIN during neuroectoderm differentiation, which was distinct from its key role in regulating the interaction of OCT4 and β-CATENIN during mesoderm differentiation.

**Figure 4.**
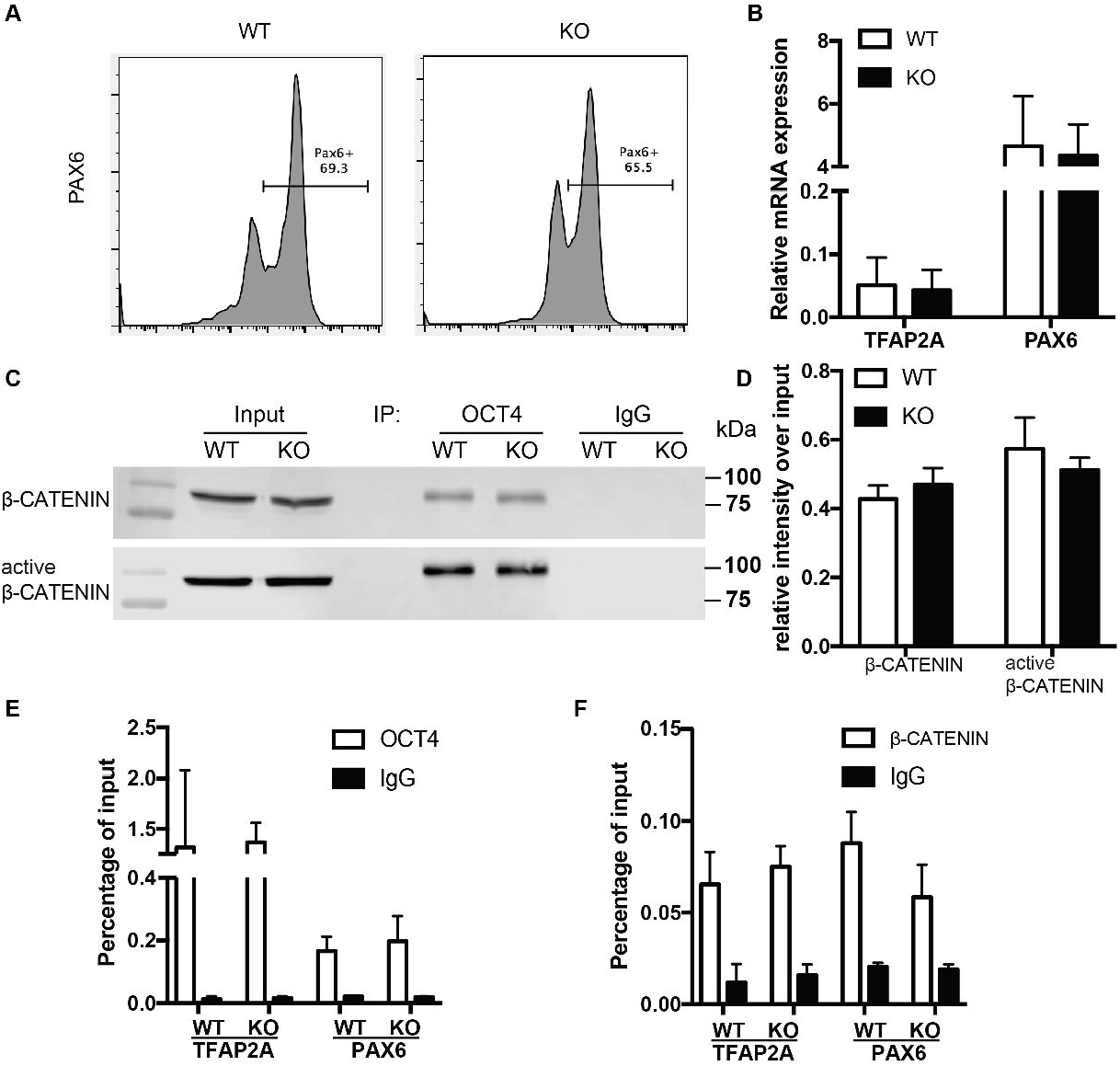
BAF250a was dispensable for neuroectoderm differentiation. (A) FACS analysis of differentiation of WT and BAF250a KO hESC to neuroectoderm using PAX6 antibody at differentiation day 12. (B) Relative mRNA expression of TFAP2A and PAX6 at day 12 differentiation. (C) Immunoprecipitation of day 2 differentiation cells after LDN193189 and SB43142 treatment using anti-OCT4 antibody, followed by Western blotting to detect β-CATENIN and active β-CATENIN. 5% of cell lysates were used as input controls. (D) Relative band intensity of precipitated β-CATENIN and active β-CATENIN in WT and BAF250 KO cells. ChIP assays with (E) Anti-OCT4 and (F) anti-β-CATENIN antibodies using day 2 differentiated cells after LDN193189 and SB43142 treatment at TFAP2A and PAX6 promoters. n=3.

### 3.4. BAF250a deletion led to prolonged S-phase in MESP1+ cells and poor differentiation potential of MESP1+ cells to ISL1+ CPCs

To further determine how the disassociation of OCT4/ β-CATENIN contributed to the later cardiac differentiation defects shown in Fig 1, we examined how the cell cycle and genes essential for cell cycle regulation were affected by BAF250a ablation, as cell cycle control is a key regulatory process for differentiation(27), and BAF250a, OCT4 and WNT/β-CATENIN are all involved in cell cycle regulation. We labeled the proliferating mesoderm cells with 5-ethynyl-2-deoxyuridine (EdU) pulse at day 3 for one hour (Fig 4A). FACS analysis was performed immediately after EdU pulse labeling and showed a higher percentage of S phase cell population and a lower G1 phase cell population in BAF250a deficient cells, indicating that BAF250a deletion led to prolonged S-phase in MESP1+ cells (Fig 4B). When we examined the EdU labeled cells at Isl1 CPC stage, we detected about 43.8%±7.4% of EdU^-^/ISL1^+^ cells and 13.0%±4.3% of EdU^+^/ISL1^+^ cells from WT hESC differentiation, but only 32.4%±5.6% of EdU^-^/ISL1^+^ cells and 1.0%±0.4% Edu^+^/ISL1^+^ cells from the differentiation of BAF250a KO hESCs (Fig 4C), suggesting that the ISL1^+^ cells are largely derived from non-proliferating mesoderm cells and BAF250a could contribute to the differentiation of ISL1^+^ cells by regulating cell cycle of mesoderm cells.

We next examined the cell cycle genes regulated by BAF250a, together with OCT4 and WNT/β-CATENIN. Expression of G1/S transition protein CCND2 and CCND3 was significantly higher in BAF250a KO cells (Fig 4D). We detected decreased recruitment of OCT4 and increased recruitment of β-CATENIN at both CCND2 and CCND3 promoters (Fig 4E,F). Since overexpression of OCT4 in somatic cell reduces cell proliferation(28) and activation of WNT/β-CATENIN leads to the proliferation of ISL1+ CPCs(29), it is likely that BAF250a regulates proper expression of CCND2 and CCND3 by modulating the recruitment of OCT4 and β-CATENIN to the promoters of proliferation driving genes. Together, our results strongly suggested that the prolonged S phase in Mesp1 cells caused by BAF250a deletion was a major contributor to defective Isl1 CPC differentiation.

## 4. Discussion

In this study, we show that BAF250a regulates cardiac lineage specification through regulating the interaction of OCT4 and β-CATENIN and their recruitment to target genes. BAF250a-mediated OCT4/ β-CATENIN interaction not only fine-tunes the MESP1 and EOMES gene expression but also regulates the cell cycle progression of mesoderm cells. We further demonstrate that BAF250a regulates G1/S transition in mesoderm cells, which is critical for the differentiation into cardiac lineage. Therefore, by modulating the interaction and recruitment of OCT4 and β-CATENIN, BAF250a guides the direct cell-type specific gene expression and the cell cycle machinery in mesodermal cells to promote cardiac lineage commitment. Importantly, our data indicate that BAF250a is a key subunit regulating SWI/SNF activity. We show that BAF250a is required for BRG1 interaction with OCT4 in mesoderm cells. In addition, the observed key role of BAF250a in regulating cardiac mesoderm differentiation appears lineage specific, as BAF250a is dispensable for the interaction of OCT4 and β-CATENIN during neuroectoderm differentiation and BAF250a deficiency does not affect neuroectoderm differentiation.

We report here that BAF250a is required for efficient interaction of OCT4 and β-CATENIN in T+ cells for proper cardiac mesodermal differentiation. Direct differentiation of cardiomyocytes from human ESCs is based on sequential stimulation and inhibition of WNT/β-CATENIN signals identified during cardiogenesis(4). Dynamic Oct4 recruitment to the genome in response to different signals is required for proper differentiation of mesoderm from hESCs(19,30) and the interaction of OCT4 and β-CATENIN is essential for mesoderm differentiation(19). In this report, we have identified that the physical interaction of OCT4 and β-CATENIN is significantly reduced in BAF250a KO cells, which leads to aberrant recruitment of Oct4 and β-CATENIN to the promoter of mesoderm genes, such as MESP1 and EOMES. OCT4/β-CATENIN has been revealed to prepattern the epigenome by regulating PRC2 recruitment to the mesoendoderm genes, whereas BAF250a/BAF complex is required for PRC2 eviction in the genome(31). In our study, we find reduced removal of H3K27me3 at MESP1 and EOMES promoters during differentiation in BAF250a KO cells. Based on these studies and our findings presented here, BAF250a-mediated OCT4/β-CATENIN interaction likely plays an important role in efficiently removing mesoderm differentiation barrier during differentiation.

Indeed, our studies show that the aberrant recruitment of Oct4 and β-CATENIN leads to altered expression of several key groups of genes that negatively affect cardiac lineage differentiation. Particularly, the expression of key mesodermal lineage gene MESP1 and EOMES is decreased. In contrast, we have identified upregulation of SOX2 and NANOG in BAF250a knockout cells during differentiation. Our finding is consistent with reported studies that the interaction of OCT4 and β-CATENIN is essential for the repression of SOX2 and activation of mesoderm genes (19,32,33). Importantly, the upregulation of neural gene such as TFAP2A we find in BAF250a KO cells could be direct consequence of OCT4 and β-CATENIN complex dissociation, because β-CATENIN induces the expression of neural genes in the absence of OCT4(19). Furthermore, we find that BAF250a deletion does not alter the interaction and the recruitment of OCT4/β-CATENIN during neuroectoderm differentiation. These results suggest that BAF250a is a key subunit defining the specificity of SWI/SNF in mesoderm cells.

Our findings on the connection of cell cycle regulation and direct differentiation in mesodermal cells highlight the importance of cell cycle regulation in direct differentiation in the multipotent cells. We find that BAF250a regulates the cell cycle of mesoderm cell through controlling the expression of cell cycle gene CCND2/3. This is consistent with previous studies that BAF complexes could act as transcriptional repressor of positive cell cycle factors in G1/S transitions(27). As CyclinD proteins have been suggested to modulate the transcriptional activity of SMAD2/3 to promote neuroectoderm differentiation of hESCs(12), it is likely that BAF250a-mediated OCT4/β-CATENIN complex could directly activate cardiac gene program while also mediating cell cycle exit. Indeed, our finding that most of Isl1+ cells are derived from non-proliferating mesoderm cells reveals a significant role of cell cycle in regulating cell differentiation. Importantly, BAF250a deletion reduces the cardiac differentiation by promoting cell proliferation, as indicated by lower cardiac differentiation potential of S stage mesoderm cells in BAF250a knockout cells. Moreover, we find that BAF250a is also required for proper recruitment of OCT4/β-CATENIN to CCND2/3 promoters in mesoderm cells. As persistence of WNT/β-CATENIN signal leads to expansion of cardiac progenitors(29) and overexpression of Oct4 leads to reduced proliferation in somatic cells(28), our results suggest that BAF250a coordinates with WNT/β-CATENIN signal to control the cell cycle exit of mesoderm cells and therefore cardiac differentiation.

In summary, we demonstrate that the physical interaction of BAF250 with Oct4 and WNT/β-CATENIN complex is critical for the cardiac differentiation by directly regulating cardiac mesodermal lineage gene expression and cell cycle phases. Our study also indicates that manipulating the cell cycle state of mesoderm cell may further enhance the cardiac lineage specification. Further studies in understanding how BAF complex interplays with developmental signals in maintaining proliferation and differentiation balance may provide key molecular mechanisms for therapies against heart disease.

**Figure 5.**
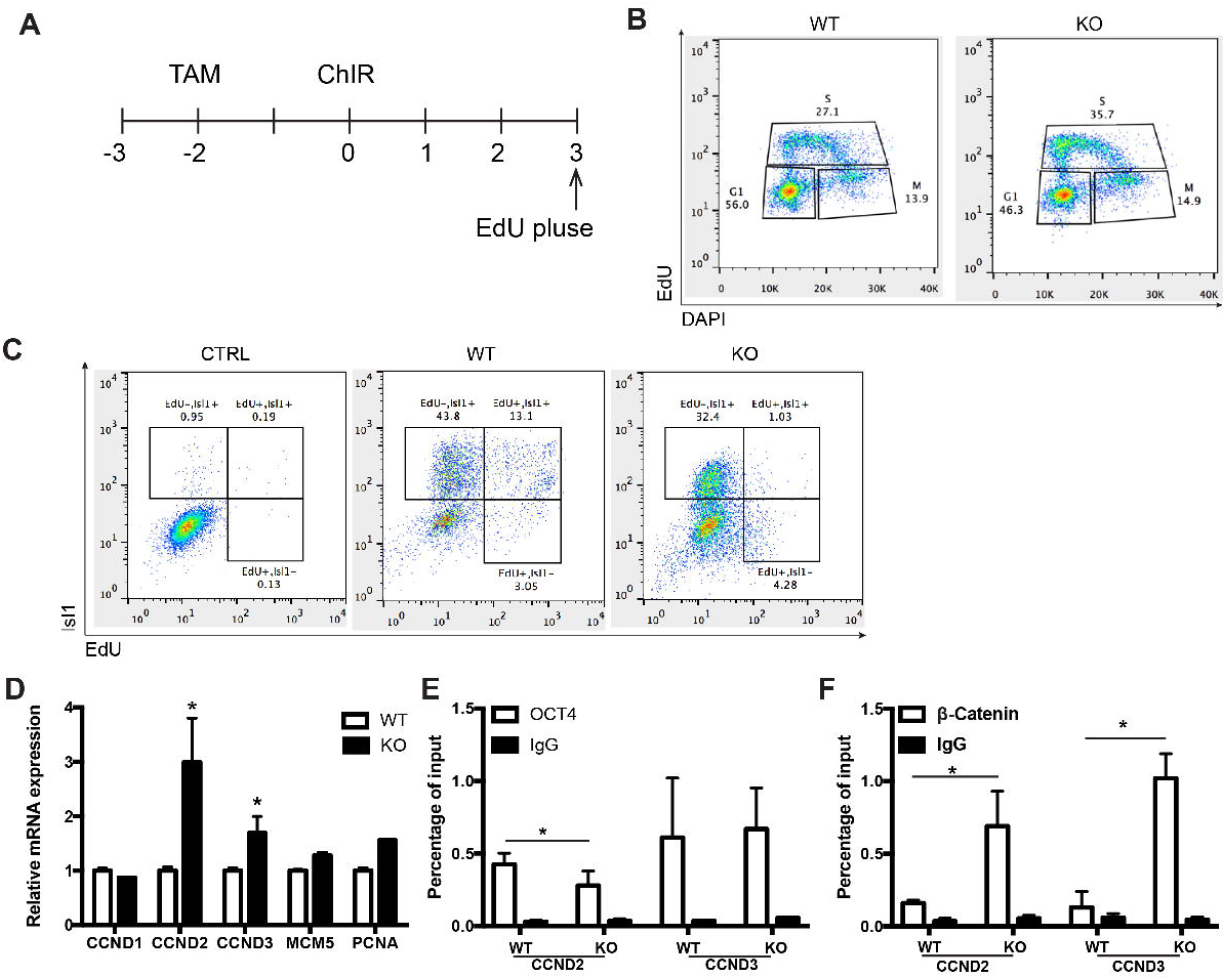
BAF250a regulated the cell cycle of MESP1 cells and promoted MESP1 cell differentiation to Isl1 CPCs. (A) Schematic diagram of EdU labeling. (B) Cell cycle analysis at differentiation day 3 with EdU and DAPI. (C) FACS analysis of Isl1+ cells at differentiation day 6 using EdU pulse labeling. (D) Expression of cell cycle genes CCND1, CCND2, CCND3, MCM5 and PCNA at differentiation day 3. ChIP assays with (E) Anti-OCT4, (F) anti-β-CATENIN antibodies using day 3 differentiated cells.

## Acknowledgments

This work was supported by the National Institutes of Health Grant HL109054 **, an Inaugural Award from the Samuel and Jean Frankel Cardiovascular Center, University of Michigan, a Pilot Award from the Joint Institute of University of Michigan Health System and Peking University Health Science Center to Z.W..

## Contributions

I.L., Y.Z. and Z.W. designed the study; I.L., T.S. and V.C. performed experiments; I.L., Y.Z. and Z.W. wrote the manuscript; Z.W. final approval of manuscript.

